# Key ecological parameters of immotile versus locomotive life Фундаментальные экологические параметры неподвижной и передвигающейся жизни

**DOI:** 10.1101/836882

**Authors:** V. G. Gorshkov, A. M. Makarieva

## Abstract

Principles of stable ecosystem organization are considered together with the role of abundant space, matter and energy in its maintenance. Life features the dichotomy of immotile (sessile, sedentary) organisms like plants, fungi, bacteria, on the one hand, versus organisms capable of active locomotion (animals) on the other. The immotile life can form a continuous live cover on the Earth’s surface. Since all available space is occupied, the immotile life does not experience an affluence of matter, energy and space itself. It turns out that this lack of abundance permits organization, on the basis of immotile organisms, of a stable ecosystem with a steady biomass. This live biomass comprises time-invariable genetic information about how to keep the environment in a stable state by controlling the degree of openness of nutrient cycles. Crucially, depending on their body size, energy and matter consumption by large animals exceed the area-specific fluxes of net primary production and its consumption in the immotile ecosystem by up to three orders of magnitude. The implication is that the herbivorous animals can meet their energy demands if and only if they move and destroy the live biomass of the immotile ecosystem. In consequence, if the immotile heterotrophs are replaced by locomotive heterotrophs, the ecosystem biomass experiences huge fluctuations and the ecosystem loses its capacity to maintain its favorable environment. From available theoretical and empirical evidence we conclude that life’s organization remains stable if the share of consumption by large animals is strictly limited, not exceeding about one per cent of ecosystem net primary production.

## 1. Introduction

Life is the most powerful process that determines the state of the environment on Earth. A vivid illustration is provided by the stores and fluxes of carbon, life’s main chemical element. With stores of organic carbon in soil and the ocean and of inorganic carbon in the atmosphere all of the order of 10^3^ GtC (1 Gt = 10^9^ t) [1,2] and with global net primary productivity of the order of 10^2^ GtC/year [3], if synthesis and decomposition get unbalanced life is able to completely perturb the environment in just about ten years. This does not however happen even on much longer timescales characterizing the lifespan of biological species (~10^6^ yr) or ecosystem (>10^7^ yr). This implies that life possesses information about the essential environmental characteristics and keeps those characteristics in a state favorable for life itself – by compensating, below a certain threshold, all unfavorable environmental fluctuations of both biotic and abiotic origin. This is the essence of the biotic regulation of the environment [4,5].

Green plants absorb solar radiation and, on this basis, with an efficiency of about one per cent synthesize organic matter that further serves as the energy source for all living organisms. Maximum power of the flux of synthesis *P* of organic matter per unit area of the Earth’s surface is of the order of *P* ~ 1 W/m^2^; it is set by the power of solar radiation and cannot be changed by life. Meanwhile the surface-specific power *J* (W/m^2^) of the flux of decomposition of organic matter – respiration – depends on the amount of live biomass per unit area. Due to the universal structure of DNA and the universal biochemical nature of living organisms the power of respiration per unit volume of living bodies is on average the same for all species of the biota and constitutes around *Q* ~ 1 kW/m^3^ [6,7], Fig. 1a. Therefore big animals with a vertical linear size *l* of the order of 1 m have the surface-specific flux of decomposition *J* = *Ql* exceeding the synthesis flux *P* by two orders of magnitude or more, Fig. 1b. Such animals are “hotspots” of grossly unbalanced synthesis and decomposition and, for this reason, they represent a potential danger to the stability of ecosystem and environment [6,8]. This fundamental property of large animals derives unambiguously from their inherent biological features but so far it has not become part of the many discussions of ecological plant-animal relationships and ecosystem stability [9–13].

**Fig. 1.**
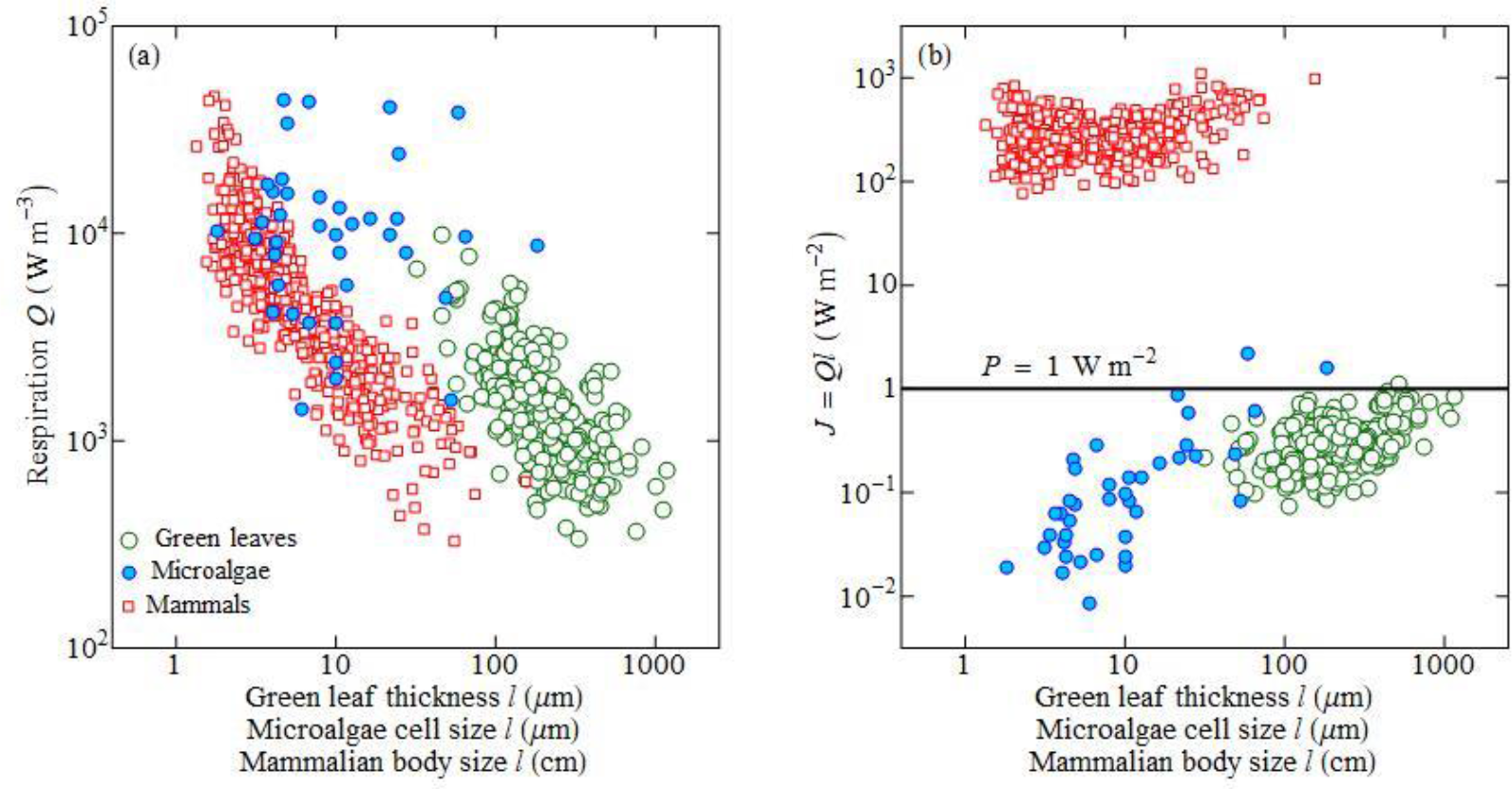
Respiration rate per unit live volume *Q* (a) and per unit projection area *J* ≡ *Ql* (b) in organisms with different linear size *l*: green leaves [14], microalgae [7] and mammals [15]. For mammals *l* ≡ (*m*/*ρ*)^1/3^, where *m* is body mass, *ρ* = 10^3^ kg m^−3^ is liquid water density. In (b) *P* = 1 W m^−2^ is a characteristic net primary productivity of the biota, see text.

Unlike big animals, small heterotrophic organisms, in particular, bacteria and fungi, with their linear sizes not exceeding *l* = *P*/*Q* ~ 1 mm, can, similar to plants, form a continuous immobile cover and thus ensure a stable flux of decomposition of organic matter balanced with its synthesis. The continuous cover of plants and microscopic heterotrophs is fundamental for ecosystem organization in two important aspects. For the first, it makes it possible for the biota to monitor environmental parameters and react to disturbances in any point of the biosphere. A simultaneous ubiquitous presence of the biotic regulation’s working mechanisms (living cells) ensures that the environmental regulation by the immotile life becomes maximally effective; it uses all the available genetic information about the necessity of the closeness of the matter cycles and compensation of any perturbations of these cycles. For the second, when life forms a continuous cover such that there is no free (unclaimed) space, matter or energy, i.e. no affluence, it is possible to efficiently stabilize the genetic program of the biotic regulation. Biotic regulation comprises ultracomplex interactions of living organisms with their environment, which also includes organisms of different species. These interactions are dictated by the genetic program written in DNA macromolecules – the species’ genome. New generations of DNA macromolecules are produced as copies of the parental generation and this copying process is prone to errors. By analogy with radioactive decay, one can introduce half-decay time for the copying process – at this time half of the copies remain identical to the original, the other half of copies carries at least one error [4,16]. The population is cleaned of such erroneous copies in the process of competitive interaction between individuals: normal individuals with error-free genomes should recognize and delete from the population individuals with erroneous genomes. This principle of maintaining information is unique to life and cannot be found in the inanimate world [17]. Decay individuals (i.e. those with erroneous genomes) that are produced by normal individuals forming the continuous cover of the immotile ecosystem (plants, bacteria, fungi) die due to the lack of free space and resources (“lack of affluence”): they just have nowhere to exist. In contrast, in populations of big animals, where no continuous cover is formed, such decay individuals may accumulate for a longer while escaping competitive interaction with normal individuals.

While in recent decades one has learnt a lot about the “anatomy” of the DNA, the information about species’ interaction with the natural environment has not been deciphered. Moreover, this information cannot be retrieved from whatsoever detailed characterization of the DNA structure. It is necessary to directly assess and study the governing principles of how life and environment interact. In this article we consider the key quantitative characteristics of living organisms that define this interaction.

## 2. Life’s universal parameters

The live plant layer is characterized by the power of photosynthesis *F* (W m^−2^) and net primary production *P* (W m^−2^), both taken per unit area of the Earth’s surface, and respiration power *Q* (W/lg_l.m._= 10^3^ W/m^3^) per unit live mass (“l.m.”) or volume taking into account that live mass density is close to the density of liquid water, 1 kg_l.m._/dm^3^ = 10^3^ kg/m^3^. These magnitudes can be expressed per unit surface area as well as per unit depth of the layer *l* (m). In particular, *J* = *Ql* ((W/m^3^) m = W/m^2^) is the respiration of live biomass layer of depth *l* per unit Earth’s surface area.

Mean respiration rate (dark respiration in plants) *Q* is of the order of one Watt per kilogram, Fig. 1a; it is a universal characteristic of life common to most taxa of the biota [6,7]:

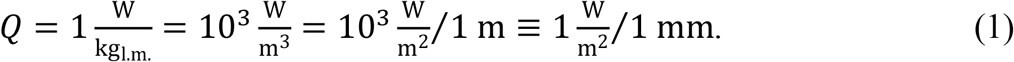

The value of *Q* (1) is universal in that sense that in any taxon independent of the mean body size of its organisms (bacteria, unicellular eukaryotes, insects, mammals, green leaves of higher plants etc.) the respiration power turns out to be of the order of one Watt per kilogram for many species. At the same time within each group of organisms *Q* may depend on body size [18]. For example, green leaves with mean thickness of 0.1 mm and mammals of body mass 1 kg respire at a rate of about 1 W/kg = 10^3^ W/m^3^, while the smaller autotrophs (unicellular microalgae) and the smaller mammals – at a higher rate of a few Watts per kilogram (a few kiloWatts per cubic meter), Fig. 1a.

Another fundamental characteristic of life is the energy content *K* per unit mass (volume) of the living body (living layer) [19], which for most species is around

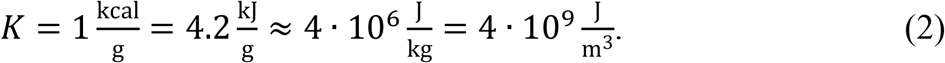

Organic carbon (C^+^) content in live biomass is of the order of 10% [7]. Energy content per unit organic carbon mass (kgC^+^) is thus about one tenth of (2),

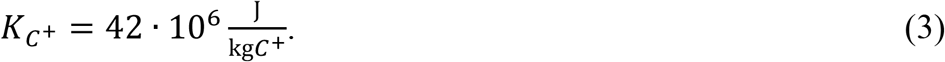

Energy power is unambiguously converted to metabolic rate with use of (2) and (3)

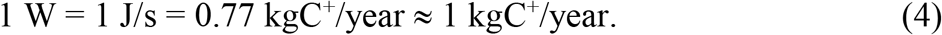

Photosynthesis (gross primary productivity) is limited from above by the flux of absorbed solar radiation and by the efficiency of converting solar energy to the energy of organic matter. A continuous vegetation layer with a minimum depth equal to one cell size already consumes all incoming solar radiation (neglecting the albedo). Thus photosynthesis on land during the vegetation period (warm months in the temperate zone and annually in the tropics) is approximately the same in all ecosystems with sufficient moistening [20,21]:

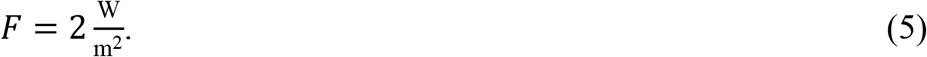

Energy produced by photosynthesis is divided between respiration and net primary production (i.e. growth and reproduction of plants). Net primary production is consumed by heterotrophs – bacteria, fungi and animals. From the law of matter conservation we have

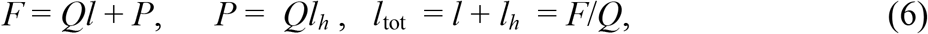

where *l* and *l*_*h*_ are the depths of the plant layer and heterotroph layer, respectively. In the temperate zone the vegetative season takes three-four months. During the rest of the year the total metabolically active layer *l*_tot_ of live plants and immotile heterotrophs can diminish down to zero.

For mature steady-state vegetation death of plant parts should be compensated by net primary production *P*. Net primary production *P* = *P*_*c*_ of mature vegetation is, in the form of dead plant matter, consumed by the immotile heterotrophs to sustain their respiration (1). According to observations [20,22–24], gross primary productivity *F* exceeds net primary productivity *P* by approximately twofold. Therefore, net primary productivity *P*_*c*_ of mature steady-state vegetation approximately coincides with the respiration of the mature vegetation layer *l*_*c*_:

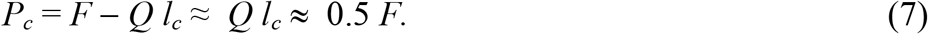

Due to this coincidence and the universal magnitude of respiration *Q* per unit depth of live layer in plants and heterotrophs, depth *l*_*h*_ of the layer of immotile heterotrophs that consume net primary productivity *P*_*c*_ = *Ql*_*h*_ ≈ *Ql*_*c*_ approximately coincides with depth *l*_*c*_ of the plant layer: *l*_*c*_ = *l*_*h*_ = 1 mm, Fig. 2:

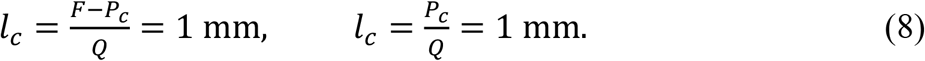

**Fig. 2.**
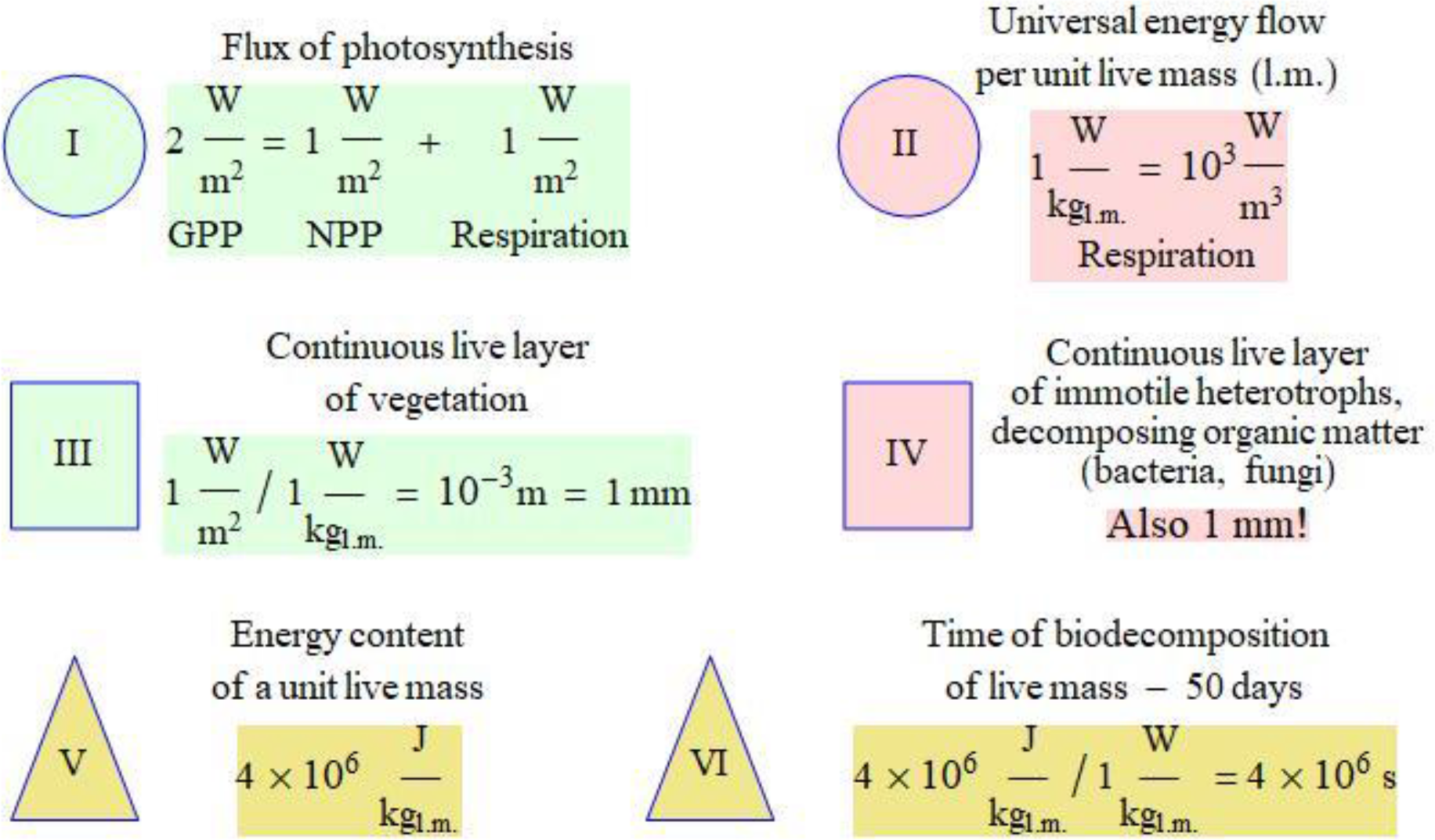
Some of life’s key numbers, see Eqs. (1)–(9); NPP and GPP are, respectively, the net and gross primary productivity.

Due to relationship (7) time *τ* = *K/Q* of decomposition of live mass (or volume) by respiration coincides with time *τ*_*p*_ = *Kl_c_/P_c_* of synthesis of layer *l*_*c*_ of live mass by plants and constitutes

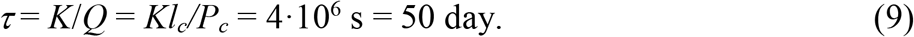

This is approximately the time of growing one harvest.

The characteristic values of biotic productivity given in Eqs. (5) and (7) and in Fig. 2 correspond to the vegetative season in terrestrial ecosystems with sufficient moistening, mostly forests [21]. The global mean net primary productivity of the biota on land is lower than that as it accounts for the unproductive territories like deserts. Global net primary production *P*_*g*_ = 54 GtC^+^/yr [3] with land area *S*_*l*_ =1.5·10^14^ m^2^ corresponds, according to (4), to net primary productivity of *P* = *P*_*g*_/*S*_*l*_ = 0.4 кгC^+^/yr/m^2^ = 0.5 W/m^2^ and *F* = 1 W/m^2^, which is half the values of (5) and (7). We also note that for forests the ratio *P*/*F* (7) can vary from 0.3 to 0.5 [20].

## 3. Ecosystem organization based on immotile organisms and biotic regulation

Life’s energy source is solar radiation. Only autotrophic organisms are capable of transforming the energy of solar photons into the energy of organic matter, which can be further used by all organisms, both autotrophs and heterotrophs, to keep alive. On land plants form the energetic basis of life. The maximum live (i.e. metabolically active) vegetation layer *l*_*c*_ = 1 mm, Fig. 3, is limited from above by the flux of solar radiation and by the universal magnitude of respiration rate per unit volume. With a mean thickness of green leaves of about *d* = 0.3 mm (Fig. 1) a live layer of vegetation *l*_*c*_ = 1 mm corresponds to a leaf area index equal to *l*_*c*_/*d* = 3.

**Fig. 3.**
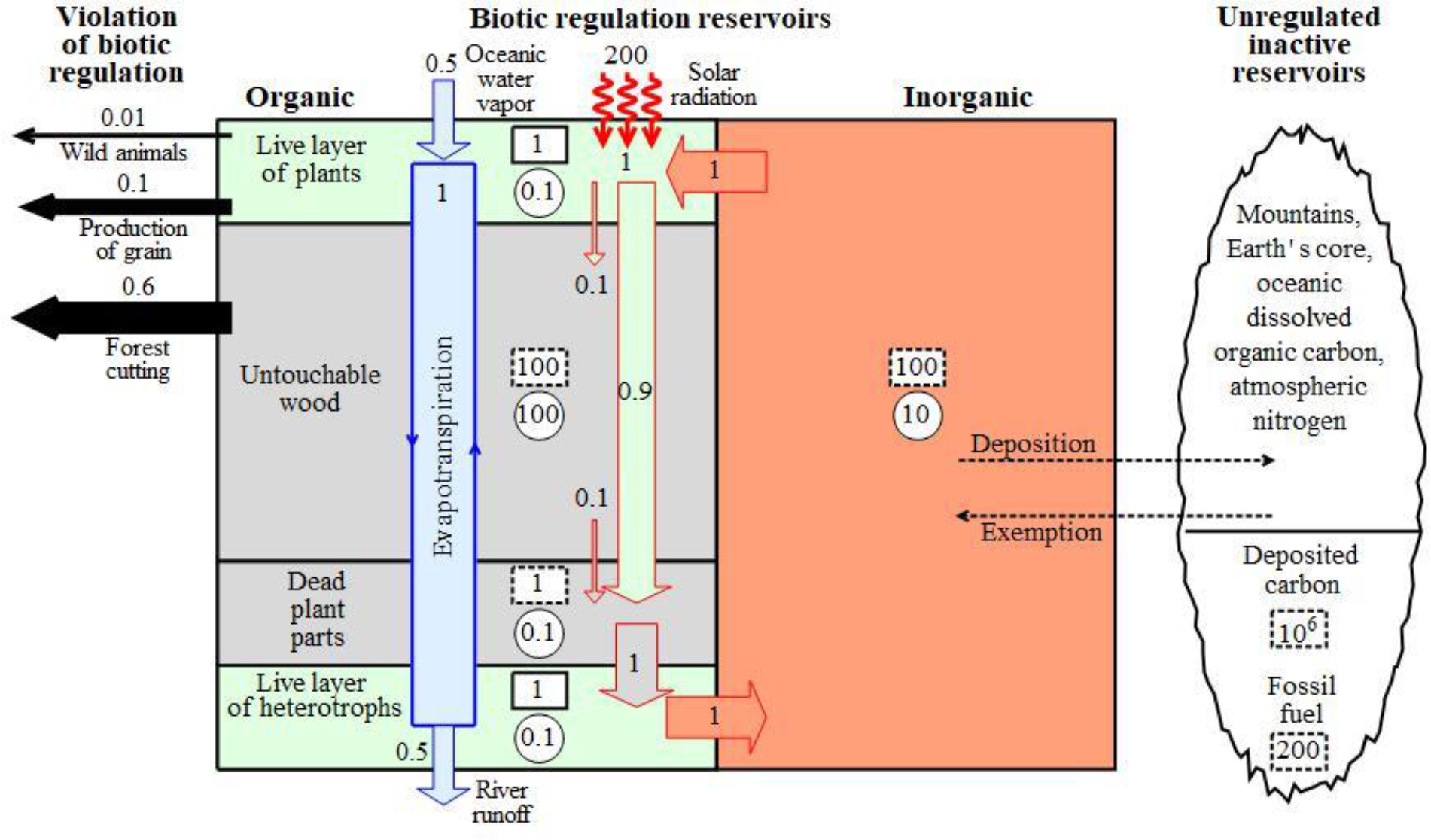
Biotic regulation of the environment by the immotile organisms of the ecosystem. Numbers in boxes are stores of carbon; metabolically active organic carbon (green) – solid boxes, metabolically inactive – dashed boxes (grey – organic, orange – inorganic) in equivalent depths of live layers (*l*, mm). Red-contoured arrows with numbers are fluxes of carbon in power units (W/m^2^), circles are carbon turnover times in years (the ratio of carbon store to carbon flux in the reservoir). Blue arrows are fluxes characterizing the water cycle (W/m^2^). Black arrows are the power fluxes (W/m^2^) of live plant destruction by various agents in the modern biosphere.

On land most part of live plants’ biomass is not green leaves but wood [25]. The wood layer in forests can exceed the layer of green leaves by two orders of magnitude reaching 100 mm (this corresponds to an organic carbon store of the order of 10 kgC^+^/m^2^), Fig. 3. The wood increment occurs at a rate of about 0.1*P*_*c*_ [22]. As the maximum plant layer *l*_*c*_ is fixed, net production of wood by plants can only take the form of such a layer of organic matter that does not respire. Indeed, compared to green leaves, the wood is metabolically inactive [25,26]. However, it performs most important structural functions including the regulation of the temperature regime within the canopy and distribution of soil moisture in soil [27,28]. High forest canopy is essential for the functioning of the biotic pump of atmospheric moisture that regulates the water cycle over forest-covered continents [29–32]. Therefore the wood of live trees is ecologically “untouchable” – it is maximally protected against being eaten and is predominantly decomposed only after the tree’s death by immotile heterotrophs (fungi, bacteria) and to a lesser degree by larger locomotive animals (earthworms, some insects and other invertebrates).

Immotile heterotrophs have a live layer approximately coinciding in depth *l*_*h*_ = 1 mm with the plant layer; the heterotroph layer cannot be increased either, Fig. 3. The heterotrophs reproduce their own live layer and decompose their own non-respiring dead organic matter resulting from dieback of live organisms. Therefore, at a fixed flux of solar radiation the total live layer *l*_*tot*_ of the immotile organisms of the ecosystem is also fixed and constitutes around *l*_*tot*_ = *l*_*c*_ + *l*_*h*_ ≈ 2 mm.

Thus, a most important property of the immotile life is that the live layer remains constant once the flux of solar radiation is fixed. The immotile heterotrophs consume net primary production in the form of dead plant parts – they do not *destroy live plants* and thus do not disturb the basic energy flux in the ecosystem – from solar radiation to organic matter. Such a constancy of the ecosystem live layer preserves the genetic information about stable closeness of the matter cycles, compensation of any external disturbances with help of directional non-random openness of the cycles and about all other ways of environmental regulation by the biota. Here the relationship *Ql* = *P*_*c*_ = *Ql*_*h*_ (7) is crucial: in the steady-state the metabolic powers (respiration) of the synthesizing (autotrophs) and decomposing (heterotrophs) blocks of the biotic regulation mechanism coincide, each constituting about one half of the maximal flux of photosynthesis *F* confined by solar radiation. These two equally powerful blocks allow the biota to react to environmental disturbances with a maximal efficiency by directionally changing the balance between synthesis and decomposition of organic matter.

All organisms of the immotile life find themselves under conditions when no free (i.e., unclaimed) space, matter or energy fluxes are available (“lack of affluence”). In such a situation all individuals with a distorted genetic program of environmental regulation (“decay individuals”) can be easily removed from the population by normal individuals of the immotile life. Indeed, due to the general principle of the lack of affluence such decay individuals don’t have space, energy or matter to live on while escaping competitive interaction with normal individuals.

Let us now consider the peculiarities of biotic regulation of the environment by the immotile organisms of the ecosystem, Fig. 3. We denote as *organic* and *inorganic biogen* a chemical element entering, respectively, organic (+) and inorganic (−) molecules used by life. Organic biogens enter the live layers of the immotile organisms in certain stoichiometric ratios [O/C/N/P]^+^. Due to the matter conservation law when the organic layers are decomposed the same ratios are preserved for the inorganic biogens in the environment. In mature steady-state ecological communities of the immotile organisms the equality

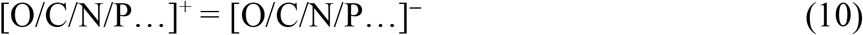

guarantees that the total environment is stationary, including the live layer *l*_*tot*_ of the immotile organisms of the ecosystem. Relationship (10) reflects the closeness of the matter cycles and the absence of the limitation by biogens: the inorganic biogens used by life are present in the environment in exactly those concentrations that they are needed.

Life transforms organic biogens into inorganic ones and vice versa, therefore both the organic and inorganic pools of biogens must be under biotic control. For example, in the case of carbon, the inorganic reservoir controlled by the biota is the atmosphere from which the plants take up carbon dioxide necessary for photosynthesis. Organic carbon reservoirs regulated by land biota are the land biota itself, Fig. 3, and the store of organic carbon in soil.

*Notes to Fig. 3*. Layer depth *l* in millimeters corresponds to a store *m* (kg/m^2^) of live biomass per 1 m^2^ of Earth’s surface in kilograms: *m* = *l*(10^−3^m)·*ρ*(10^3^ kg/m^3^) = *l* (mm)(kg/m^2^). The energy store of live mass per 1 m^2^ is obtained by multiplying mass per unit area *m* = *l*(mm)(kg/m^2^) by energy content *K* = 4·10^6^ J/kg of live mass. Turnover time *τ* = *Km*/*Q* ≈ 0.1 year for *l* = 1 mm. Carbon store and flux in the inorganic reservoir originally expressed in kgC^−^/m^2^ и kgC^−^/m^2^/year are expressed in equivalent depths of live layer (after the inorganic carbon is consumed by plants) using the relationships 1 kgC^−^ = 1 kgC^+^ (stoichiometry), 0.1 kgC^+^= 1 kg live mass and 1 W = 0.77 kgC^+^/year, see (4). Immotile heterotrophs consume dead plant parts at a rate equal to their production by plants, such that this metabolically inactive layer (litter) does not exceed the live plant layer *l* ≈ 1 mm.

The genetic program of biotic regulation is aimed at sustaining the internal milieu of the live layer, in particular, at preserving its depth (i.e. the steady-state biomass). If the cumulative amount of biogens in both organic and inorganic reservoirs were constant, then a random excess of biogens in the inorganic reservoir (e.g. a carbon dioxide surplus in the atmosphere) would mean an equal shortage of biogens in the organic reservoir (e.g., carbon loss from soil). Biogens in different reservoirs are mixed in a dramatically different way – for example, a local but significant loss of organic carbon from soil by respiration causes a minor but global increase of atmospheric carbon dioxide. In such a situation the double disturbance of biogen concentrations simultaneously in the organic and inorganic reservoirs would make it impossible for the biota to restore the environment on both global and local scales rendering efficient biotic regulation impossible.

On the other hand, external disturbances can lead to a random change of biogen content in one of the two reservoirs (organic or inorganic). If biotic regulation aimed to preserve only the internal milieu of the live layer (i.e. it stabilized the organic reservoir only), then such random external disturbances would disrupt the constancy of mass of the active reservoir of the inorganic biogen. Then the external environment were changing in a random direction under the influence of external disturbances. In the result, it would drift away from the optimal conditions and a further stabilization of the maximum possible mass of the active layer of the immotile ecosystem would become impossible. In the end, life’s stationarity and stability would become impossible as well.

Therefore, for the biotic regulation to be effective, a third reservoir, either organic or inorganic, is needed, Fig. 3, which will play the role of a buffer from which the biota could replenish the lacking biogens and where it could dispose of the excessive biogens. If the amount of biogens in the active reservoirs becomes too high, a certain part of biogens can be removed from the environment and deposited in the inactive reservoirs, Fig. 3.

A major feature of the inactive reservoir is that it should be inert with respect to the biota functioning. For example, the inactive reservoir of nitrogen is the atmosphere – the atmospheric nitrogen is well mixed globally, it is not directly used by plants nor it is a greenhouse gas. Furthermore, its store is so large compared to the soil nitrogen used by the biota that no changes in the active nitrogen reservoir regulated by the biota (soil) would ever lead to a considerable change of the atmospheric concentration of nitrogen.

Another example of an inactive reservoir is provided by the organic matter that is resistant to biotic decomposition or removed from the biosphere altogether. Generally, metabolically inactive organic matter can exist in multiple forms and be part of both regulated and inactive reservoirs. The “untouchable” wood of live forest trees that is protected from heterotrophs and covered by live metabolically active cambium (bark), Fig. 3, represents a biotically regulated reservoir. Long-lived fractions of the oceanic dissolved organic carbon, humus and soil, bog mire that are not available for decomposition by heterotrophs represent inactive reservoirs, Fig. 3.

Finally, dead matter produced by the biota with a violation of relationship (10), i.e. in the form of hydrocarbonates, hydrocarbons etc. can be removed from the regulated environment and deposited in dispersed inorganic (like mollusk shells) or organic matter in sediments. Such biotic sedimentation of organic carbon during the Phanerozoi has prevented a catastrophic accumulation in the atmosphere of carbon dioxide filtered from the Earth’s core [33]. The spatially concentrated part of this inactive reservoir of organic carbon is consumed by modern civilization in the form of fossil fuel, Fig. 3.

We emphasize once again that with all those inactive organic reservoirs present the immotile life features live layers of the order of *l*_*tot*_ = 2 mm for plants and heterotrophs, while in the regulated environment relationship (10) is fulfilled.

## 4. Locomotive animals in the immotile ecosystem

An increase in body size in evolutionary lineages (the so-called Cope’s rule, see [34]) is possible when larger species are more competitive and can force out the smaller ones from their favorable environment. Bodies of large animals can reach more than one meter in linear size, which corresponds to 10^3^ layers of the immotile life. Consequently, the respiration rate per unit area of the animal body projection on the ground surface can reach *J* = 10^3^ W/m^2^. It is a thousand times higher than the characteristic net primary productivity of plants, *J* = *Ql* ≫ *P*, Fig. 1b.

A most important ecological consequence of this relationship is that large animals cannot form a continuous cover feeding on the *production* of the immotile plants in the form of their dead parts. Large animals must move continuously across the Earth’s surface and consume live plant biomass – thus destroying live plants and introducing disturbances into the fluxes of organic matter synthesis. Furthermore, as far as the bodies of large animals do not form a continuous cover, these animals exist under conditions of abundant free space and food. This creates a possibility of an explosion-like increase in population density of big animals with an associated complete destruction of plant biomass [6,8].

The condition for existence of locomotive animals consuming live biomass of immotile plants is determined by the law of energy conservation. Let us define the linear size *l*_*a*_ of the animal to its live body mass as *l*_*a*_ ≡ (*M*/*ρ*)^1/3^, where *M* and *ρ* are the mass and density of the living animal body. Let us consider layer with depth *l* of live plant biomass distributed in space across height *H* (for example, *H* can represent the height of forest canopy or the depth of the euphotic layer in the ocean), Fig. 4. Moving with a mean daily speed *u* the animal scans per unit time a band of area *u l*_*a*_ (m^2^/s) and consumes within this band a share *γ* of the immotile live plant biomass. The rate of energy consumption by the animal is then given by *Kεγ u l l*_*a*_, where the energy content *K* of live matter is defined in (2), *ε* ≡ *l*/*H* and *εγ l l*_*a*_ *u* is the volume of plant biomass consumed by the animal per unit time, Fig. 4.

**Fig. 4.**
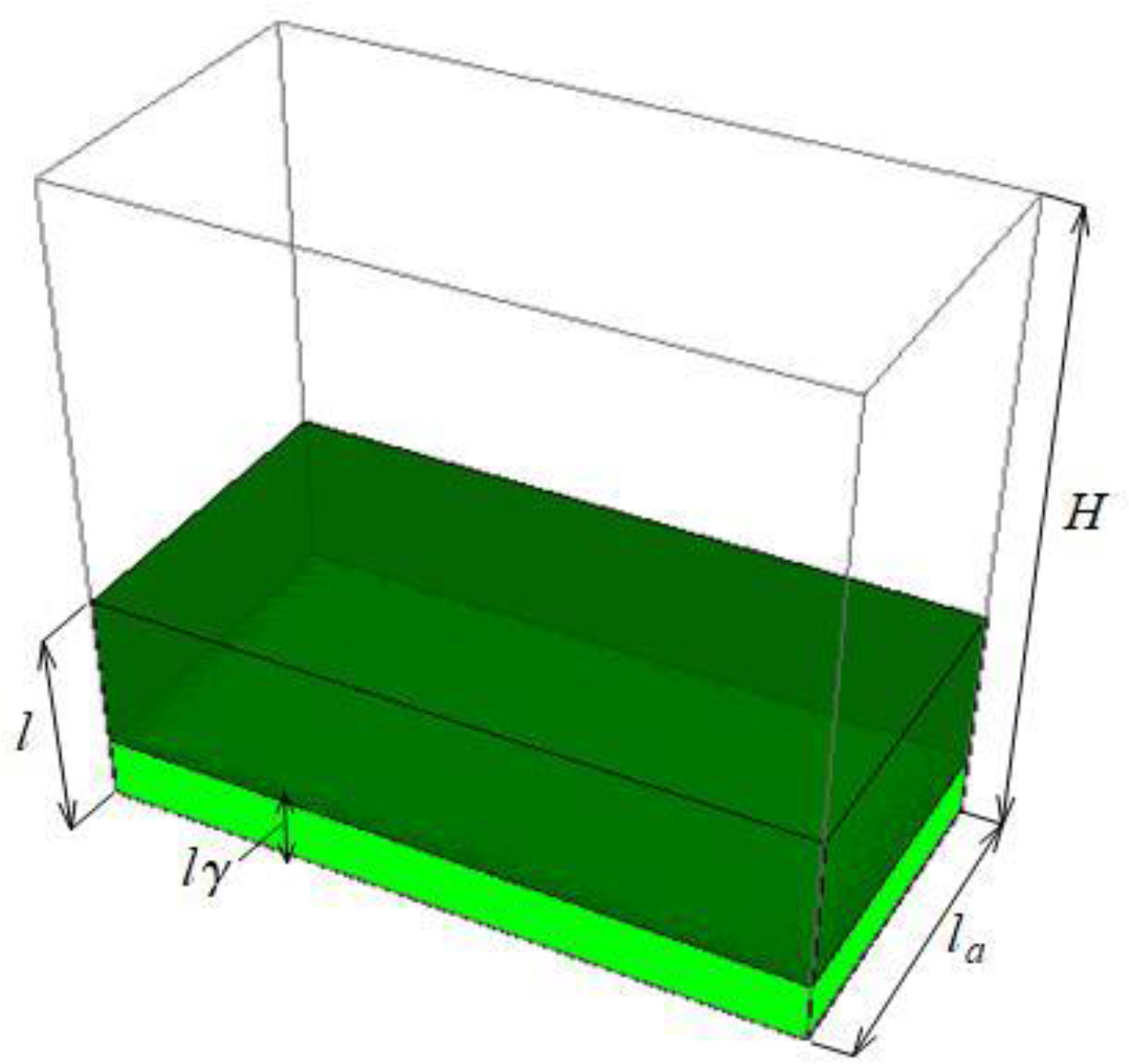
Parameters governing live plant biomass consumption by a locomotive herbivorous animal, see Eq. (11): *l* = 1 mm is the depth of live plant layer; *l*_*a*_ is the linear body size of the animal; *γ* ≤ 1 is the share of the available part of the plant layer consumed by the animal; *ε* ≡ *l*/*H*, *H* is the height of the ecosystem along which the live plant layer is distributed (for simplicity in the figure the layer is shown as a monolith but in reality it consists of leaves etc. distributed in the vertical along *H*).

The power provided by food consumption should exceed the mean daily respiration power of the animal [8]:

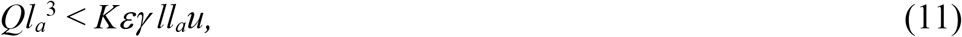

The difference between the power of food consumption by the animal and its respiration power is equal to the kinetic power of animal locomotion. While all energy consumed by the immotile life undergoes dissipation to heat within the continuous live layer, the kinetic energy of the locomotive animal undergoes dissipation to thermal radiation outside the animal body in the external environment.

The mean daily speed *u* of animal locomotion is determined by the animal ability to consume food; it can be unambiguously calculated from the data on the daily mean energetic cost of locomotion (an analogy of petrol amount spent per 100 km driving). According to numerous measurements, the daily mean speed of locomotion on land constitutes on average *u* = 0.3 m/s and does not depend on body size in mammals and birds (see Fig. 1 in [35]). For reptiles this speed is an order of magnitude lower and constitutes *u* ~ 0.03 m/s (see Fig. 8 in [36], Fig. 5.7.1 in [4]).

**Fig. 5.**
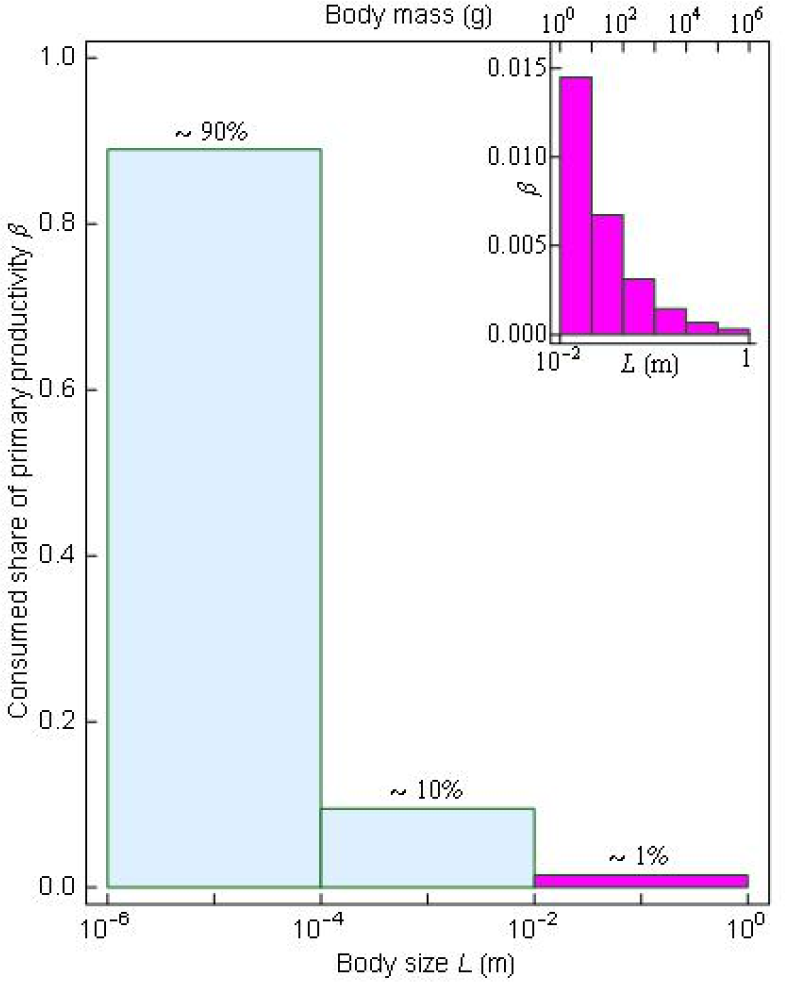
Distribution consumption of net primary production across heterotrophs with different body sizes [6,37]: 90% of plant production is consumed by the smallest heterotrophic organisms (bacteria and fungi); invertebrates (the smallest locomotive animals) consume about 10% of net primary production, vertebrates consume around 1%. The red diagram shows consumption by herbivorous mammals and birds in boreal forests [37].

Using the definition *τ* ≡ *K*/*Q* (9) we can find from Eq. (11) the maximum body size of herbivorous animals in different regimes of plant biomass consumption:

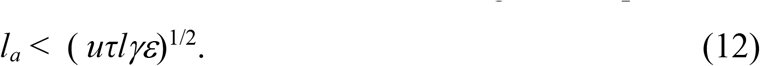

Obviously, the maximum possible size corresponds to the regime when all the dominant vegetation is fully destroyed (tree felling exemplified by elephants, beavers or forest industry in the modern times; in the past, possibly, the largest mammals like the balochiterium, mammoths and others). Tree felling and consumption of all live biomass at the ground corresponds to *γ* = *ε* = 1. In the result from Eq. (12) we obtain the upper limit on animal body size *l*_*a*_ < (*u τ l*)^1/2^ = 40 m, which includes all animals ever existed. We note that tree felling and the removal of tree canopy (clear-cutting) totally destroys the biotic regulation of the water cycle (biotic pump of atmospheric moisture) [29].

Let us further consider total consumption (*γ* = 1) of the biomass of non-woody plants of the lower stratum of the ecosystem (grasses, mosses, lichens) in a forest ecosystem where such non-woody plants constitute a minor part^1^ *ε* = *l*/*H* ~ 10^−3^of total live biomass. In this case for *u* = 0.3 m/s we obtain from (12):

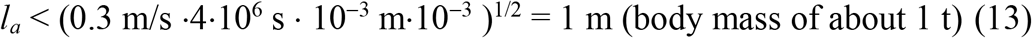

According to available measurements, the maximum values of *u* = 0.8 m/s are recorded for the donkey and African elephant [4,8]. Assuming that the same speed characterized mammoths and possibly other large extinct mammals we have from (12)

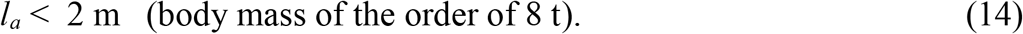

These estimates show that the largest mammals in closed canopy forest ecosystems exist on the verge of their energetic capabilities. In other words, the existing plant resources available to the moving animal are barely sufficient to cover the energetic needs of the animal.

Conditions (11)-(12) mean that the animal moving across its feeding territory with a daily mean speed *u* and consuming share *γ* of the available plant biomass layer of depth *l* obtains enough energy to cover its metabolic needs. However, these conditions do not confine the share of net primary productivity of the ecosystem consumed by the animals of a given size. This share depends on the population density of the animals or on its inverse value – the feeding area per individual.

Let us introduce the linear size of individual feeding territory of an animal with linear body size *l*_*a*_ such that the feeding area is equal to *S* ≡ *L l*_*a*_ (m^2^). We also introduce a characteristic time during which the animal scans all its territory, *τ*_*a*_ ≡ *L*/*u*. В этом случае доля *β* потребления чистой первичной продукции *P* животным составляет

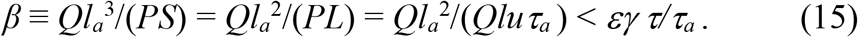

As one can see from (15), the maximum value of *β* is reached when the animal covers its feeding territory in time *τ*_*a*_ equal to the turnover time *τ*(9) of plant biomass. With *τ*_*a*_ = *τ* and *εγ* = 1 we have *β* < 1, which means that the animal can claim all ecosystem productivity.

In this case for an animal with *l*_*a*_ = 1 m and body mass 1 t the individual feeding territory will constitute *S* = *l*_*a*_*uτ* = 1 km^2^, and biomass will be 1 t/km^2^. This coincides, in its order of magnitude, with the biomass of large mammals in modern protected national parks in savannas which, however, are not stable ecosystems and suffer from overgrazing [37]. There is a view that similar biomass values were characteristic on the so called mammoth steppes [13].

The ecological problem of ecosystem instability in the presence of large herbivorous animals is manifested, among other things, in the fact that the turnover time *τ*(9) of plant biomass, about two months, is significantly shorter than the lifespan time *τ*_*l*_ of the majority of terrestrial plants,

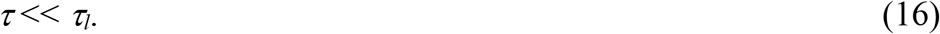

Indeed, herbivores can consume all plant biomass at a rate equal to its regrowth only during the plant’s lifetime. For example, an animal can consume the biomass of perennial herbs preventing normal formation of the seed bank and normal reproduction. The animal can also consume tree seedlings and saplings as long as the mature tree produces seeds for them. When condition (16) holds, animal biomass can be accumulating for a long time *τ*_*l*_, which in case of trees is of the order of a few hundred years. During this time the population of large animals can increase to a point when the animals will consume all young seedlings produced by mature trees. Then, as soon as the old tree dies, the regeneration of biomass and ecosystem recovery become impossible, the environment degrades in an uncontrollable manner, the production of edible biomass sharply shrinks and the population of large animals goes extinct together with the original ecosystem. In most detailed way this process is described and studied for savannas, where it is known that an artificial increase of herbivore population density causes the ecosystem to go from the state of a woody savanna to desert [9].

However, in a broader theoretical context the problem of ecosystem instability due to the presence of large herbivores has not received much attention in the ecological literature. One of the reasons is that the mathematical equations population dynamics, in particular, the various modifications of Lotka-Volterra equations, do not presume a scenario of the population going extinct. This problem was termed the *atto-fox problem* (atto- corresponds to multiplication by 10^−18^) [38,39]. In these mathematical models even if the population density drops to an infinitely small value – e.g. to 10^−18^ foxes per square kilometer as in [39] – such a population can still recover to realistic values of population numbers. Thus the extinction scenario simply does not exist; it must be manually prescribed. If extinction is not manually enforced, the population will exist eternally oscillating between infinitely small and observable population density values. It is also possible to manually exclude scenarios when the population density becomes too low. For example, in the mammoth steppe model once the population density of large herbivores decreased below one individual per thousand square kilometers, it was automatically increased up to this value [13]. (It is pertinent to note that many large animals have individual home ranges of the order of a thousand square kilometers [40], thus for such animals population density 10^−3^ ind/km^2^ is normal rather than extraordinarily low.) Obviously, ecosystem stability in such models cannot be investigated.

According to the paleontological evidence, an inherent feature of the ecological community of the mammoth steppe was its spatial and temporal instability [41]. The megafauna of the mammoth steppe underwent repeated regional extinctions after which it recovered at the expense of migrations from distant refugia. (During the characteristic time *τ*(9) of the depletion of the organic matter energy content, 50 days, an animal with a mean daily speed of *u* = 0.3 m/s can travel over 1300 km.) Such spatial and temporal dynamics is consistent with the proposition that due to its powerful destabilizing impact on the regional environment and climate the megafauna disrupted conditions favorable for its own existence. Then the megafauna got extinct and could re-colonize the same territory only after the ecosystem has recovered in the course of succession in the megafauna’s absence.

Despite the paucity and incompleteness of the data about environmental characteristics of the mammoth steppe possible mechanisms of how the megafauna could destabilize the environment can be easily outlined. For the first, the grass layer sustained by the megafauna is not able to efficiently store moisture in soil as compared to the forest (or tundra in higher latitudes); thus the grass ecosystem cannot function as the biotic pump of atmospheric moisture [42]. Besides soil moisture control, big herbivores can change the vegetation species composition such that grasses using C_4_ photosynthesis might begin to dominate (for example, some *Artemisia* species [43,44]), which have a lower transpiration rate per unit carbon mass fixed than the C_3_ plants. Low evaporation reduces the intensity of the biotic pump. In the absence of the biotic pump precipitation on land is scarce and irregular; droughts and floods are frequent that may cause a decline in primary productivity and disruption of the food resources.

For the second, according to the available data from long-term experiments, an elevated fertilization-caused productivity of high latitude ecosystem (in particular, tundra) leads to a rapid depletion of soil organic carbon [45]. This implies that the high productivity of grasslands that is necessary to feed dense populations of big animals could be of transient nature and accompanied by soil degradation, depletion of food resources and extinction of large animals after which the successional recovery of the ecosystem began. Notably, neither the biotic pump nor soil degradation effects have been considered in mammoth steppe studies [13].

## 5. Large animals and ecosystem stability

A fundamental ecological problem is the question about the share *β* of primary production which large locomotive animals can consume without undermining ecosystem stability. As discussed in Section 2, immotile heterotrophs can meet their energetic needs by consuming only dead plant parts without disturbing live biomass. Theoretically in such a regime the fluctuations of synthesis and decomposition could be reduced down to zero. However, all living organisms are mortal. Death of plants inevitably introduces fluctuations into the ecosystem energy fluxes. Fall of old trees and associated plants and the subsequent recovery of the biota within thus formed gaps is a key process that determines the organization of forest ecosystem [12].

After total destruction of the steady-state mature vegetation (death of an old tree) and with its subsequent recovery depth *l* of the plant layer grows from *l* = 0 (seeds) to its maximum value *l* = *l*_*c*_. Gross and net primary productivity *F* and *P* first decrease down to zero at *l* = 0, while the immotile heterotrophs (bacteria and fungi) continue to respire. This imbalance reduces the amount of organic matter in the disturbance area. As the vegetation begins to recover, the gross primary productivity reaches its maximum at a certain point; with a further growth of *l* and, consequently, dark respiration *Ql*, net primary productivity *P* (6) then decreases from *F* to its initital steady-state value *P*_*c*_:

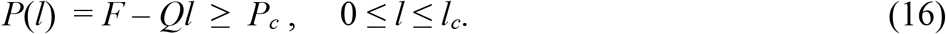

Tree death leads first to a dominance of decomposition over synthesis with an organic matter loss. Then, as the gross primary productivity has recovered, synthesis dominates over decomposition such that the ecosystem biomass regrows, while the power of heterotrophic respiration is lower than its steady-state maximum value (about half of gross productivity). During all this time the efficiency of the biotic regulation of the environment is reduced since all ecosystem resources are directed at self-recovery.

Thus in forest ecosystem the spatial and temporal fluctuations of synthesis and decomposition are set by the population dynamics of trees. Maximum environmental stability corresponds to a stationary uneven-aged population of forest trees with a fully realized gap dynamics. Consumption of live plant biomass by large animals should not perturb these patterns. In other words, in stable ecological communities plants should be protected against consumption by herbivores in such a manner that the fluctuations introduced by large animals do not exceed fluctuations dictated by the population dynamics of plants themselves.

The condition that fluctuations introduced by herbivores don’t grow with animal body size determines that share *β* of consumption of primary productivity by animals of a given size declines inversely proportionally to the linear body size, while total consumption by all vertebrates (body size greater than 1 cm) should be of the order of 1% of net primary production of forest ecosystem [6,8,37]. This theoretical result is confirmed by the analysis of empirical data on population densities of mammals and birds in boreal forest ecosystems [37], Fig. 5.

In tropical forests the share of plant production allocated to big animals (fruits and seeds) also turns out to be of the order of 1% of primary productivity. With a typical primary productivity of tropical forests around 2 kgC^+^/m^2^/yr [21], production of fruits and seeds does not exceed 0.4 tC^+^/ha/yr [46,47]. Even if all these fruits and seeds were completely eaten up by large animals, their share of consumption of net primary production would not rise above 2%.

So far debates concerning the relatively recent disappearance of large vertebrates from different continents are mostly focused around two hypotheses: climatic change and human impact (reviewed in [48]). The fundamental instability associated with potential destruction of vegetation cover by large herbivores followed by unfavorable climatic change is not taken into account. From the biotic regulation viewpoint, large herbivores could not have destroyed all life on land solely because their destabilizing impact was spatially limited: they appeared in the course of evolution, destroyed vegetation cover and went extinct together with the species constituting their food base, or without them, in relatively small regions. Then the destroyed regions were colonized by ecosystems without such disturbing agents like destabilizing large animals. Possibly, elephants, giraffes, rhinoceros and other big African species contributed to the transformation of tropical forests to savannas and deserts over larger areas. While savanna is undeniably an ancient ecosystem [49], this unstable transition between forests and deserts could have occupied a much smaller area than it does now in the absence of big herbivores. Bisons could be related to North American deserts. Australian desertification might have also been related to herbivores’ activity even before the first humans arrived. Long-term stability of the biotic regulation and matter cycles on land existed in those land regions where locomotive animals lived along rivers, lakes and seashores, i.e. on territories with a cumulative area of about 1% of the forest ecosystem area, where animal disturbances of plant cover coincided in the order of magnitude with the geophysical disturbances.

Let us emphasize a principal difference in the organization of terrestrial versus oceanic ecosystems. In the ocean photosynthesis is performed by microscopic phytoplankton which has a live biomass an order of magnitude smaller than the green leaves on land (Table 1). For unicellular organisms their lifetime *τ*_*l*_ and time of biomass synthesis *τ* coincide, therefore a prolonged increase in population numbers of herbivorous animals feeding on phytoplankton is not possible, cf. Eq. (16). Furthermore, depth *H* ~ 200 m of the euphotic layer where phytoplankton is distributed exceeds forest canopy height by almost an order of magnitude. According to Eq. (12), the small depth *l* of the live photosynthetic layer in the ocean and its distribution over a large depth *H* make it impossible for large herbivores to meet their energy demands by feeding on oceanic phytoplankton and disturbing its functioning. Large herbivores are absent in the ocean, while large animals in general consume a negligible share of ecosystem productivity (see, e.g., [50]), such that biotic regulation in the ocean is stably preserved.

**Table 1.** Productivity and biomass on land and in the ocean

|  | Live biomass<br>(GtC<sup>+</sup>) | NPP<br>(GtC<sup>+</sup>/yr) | Biomass/NPP<br>(yr) | Respiration<br>(W/kg) |
| --- | --- | --- | --- | --- |
| Ocean (phytoplankton) | 1.3 [25] | 49 [3] | 0.027 | 8.8 [7] |
| Land (green leaves) | 15 [25] | 56 [3] | 0.27 | 1.2 [7] |

In contrast to the ocean, terrestrial ecosystems cannot exist without rain and rivers. Rain and rivers are sustained by the flux of atmospheric moisture from the ocean to land. This flux is maintained by undisturbed mature forests representing continuous layer of high trees [29,42]. Such an ecosystem structure necessitates large stores of live plant biomass that can be potentially destroyed. Thus namely land ecosystems are inherently vulnerable to disruption by large animals including humans.

## 6. Discussion: Necessary and sufficient conditions for the biotic regulation of the environment

We have considered the major energetic characteristics of the ecosystem related to synthesis and decomposition of organic matter and body size of living organisms. Unlike plants and microscopic heterotrophs large organisms cannot form a continuous cover as they consume, per unit area, a hundreds of times higher energy flux than the vegetation cover produces, Fig. 1b. For this reason large animals do stand “elbow to elbow” but exist under conditions of abundant space, matter and energy.

Due to this abundance of free space, at any given moment of time the animal receives information only about a tiny part of its feeding territory. Thus it can only regulate the internal milieu of its own body and the tiny spot it instantly occupies. Therefore if the immotile heterotrophs (bacteria, fungi) are replaced by large locomotive animals consuming all net primary productivity, the biotic regulation of the environment becomes impossible.

A second important consequence of abundance is the reduced efficiency of competitive interaction. Lack of abundance (free space) in the immotile organisms forming a continuous cover guarantees the simplest way of doing away with the decay individuals (those with the genetic program of biotic regulation distorted by mutations). Such individuals are just deprived of free space and cannot escape competitive interaction with normal individuals. In locomotive animals this process of clearing the population from decay genetic information becomes much more complicated. Under conditions of naturally abundant space and edible biomass the decay individuals can exist escaping competitive interaction with normal individuals. To remedy the situation, many animal species have a genetic program of intense social interaction manifested most commonly as long migrations. The biological meaning of this program is to liquidate the abundance of space by formation of nearly continuous dense flat herds (of deer, antelopes, elephants) on the Earth’s surface or spherical (or spatially distributed in another strictly specified manner) animal crowds in the air or in the water (fish schools in the ocean, flocks of starlings, geometric wedges of geese, “military parades” of loons etc.). In such a crowded condition the abundance of space disappears. In the result, the decay individuals are more easily identified and forced out from the population by the normal individuals of the same species as well as by predators. Predators increase their population densities as well around the prey herds and also diminish the abundance of space.

Biotic regulation of the environment can be performed by ecosystems consisting of 1) immotile plants producing organic matter and 2) immotile heterotrophs decomposing this organic matter into inorganic compounds, which 3) both form a live continuous layer on the Earth’s surface of the immotile ecosystem that functions in the absence of abundant free space, matter and energy with 4) the quota of consumption of plant production by large animals being limited within about 1% of net primary productivity of the steady-state mature ecosystem, Fig. 5. Besides, it is neccessary 5) that the genomes of plants and immotile heterotrophs contain information about the rigid correlation of their functioning which ensures that in the absence of external disturbances the matter cycles are closed, while in the presence of external disturbances the matter cycles open in a non-random compensatory manner and 6) that there exists an inert inactive reservoir of organic or inorganic biogens, Fig. 3. These six conditions are necessary and sufficient for the biotic regulation of the environment to operate.

If at least one of these conditions is not fulfilled, for example, if primary producers and heterotrophs are represented by an artificial assortment of alien species, or if the inactive reservoir of biogens is absent, the ultra-complex genetic program of biotic regulation does not work and the environmental regulation is impossible.

A most important condition is the immobility of plants which enables plants to cover all land surface. The continuous plant cover does away with the abundance of space. Plants being immotile is a phenomenon of physical rather than biological origin. The energy for organic matter synthesis comes to plants in the form of solar radiation, which consists of particles – photons. Photons are the only known observable particles with zero mass.

Only objects with a non-zero mass can accumulate. The energy of photons that have zero mass cannot accumulate on the Earth’s surface. Thus plants have to limit their energy consumption to the flux of solar photons per unit area of the surface occupied by the plant. Once all surface has been occupied, a continuous plant cover of a maximum depth formed, and all flux of solar photons claimed, plants cannot move since there is neither free space nor available energy for such movement. If the Sun were sending to the Earth some energy-rich organic matter with non-zero mass, for example, a flux of organic substances on which plants could feed, then the depth of the live layer could have locally increased (in the locations where these substances accumulated). In other places, by energy conservation, the live layer would diminish or break yielding free space and creating an opportunity for plants to move. In the result, the continuous plant cover would disappear producing abundant free space, matter and energy. The biotic regulation of the environment and life itself could not exist.

The existence of immotile heterotrophs decomposing the organic plant matter on land is, in contrast to plant immobility, of biological rather than physical nature. It is an evolutionary discovery of the land biota – that immotile heterotrophs like fungi can exist in the form of a continuous live layer claiming with their living bodies all the ground surface. In the oceans fungi are practically absent [25]. Continuous plant cover and continuous cover of heterotrophs functionally correlated to the plants ensure an efficient monitoring of environmental information across the entire land surface, such that an efficient program of biotic regulation of the environment can be written into the genetic program of the immotile organisms.

Energy consumption by the immotile life and its spending on respiration, production of organic matter and metabolism does not perturb the constant depths of the live layers of autotrophic and heterotrophic organisms. Therefore at any moment of time all genetic information about environmental regulation is available for life to perform an efficient control of environmental conditions. Being capable of changing the stores of organic and inorganic biogens by a hundred per cent in just a few years the biota nevertheless keeps these stores in a constantly favorable conditions under any external disturbances for hundreds of millions of years.

Large herbivorous animals cannot live on the fluxes of matter and energy generated by the immotile organisms without destroying their live layers. Large animals can only live by destroying the live biomass of the immotile organisms depriving the ecosystem of the ability to regulate the environment under the forming conditions of abundant space, energy and matter. Thus life and biotic regulation are only possible when energy consumption by large animals is limited to about one per cent of ecosystem net primary productivity for all vertebrate animals combined, Fig. 5. Large animals existing within this ecological corridor do not pose a threat to ecosystem stability and may even perform useful functions, e.g. spreading seeds. Humans as a large animal species have currently exceeded their ecological quota in the global scale by almost two orders of magnitude consuming about 10% of global net primary productivity [4]. As a consequence, terrestrial ecosystems have globally degraded and regional and global climate destabilization followed. Protection and restoration of the remaining sustainable ecosystems on land should be a major goal in the global change context [51].

## Footnotes

1 Note that one and the same share *ε* = 10^−3^ of available plant biomass can characterize different spatial configurations of plant biomass distribution in the ecological community. For example, if the animal feeds on openings resulting from the dieback of large trees, and such openings constitute 10^−2^ of forest area, and on each such plot the animal eats up about one tenth of available plant biomass, we get the proportion *ε* = 10^−3^.

